# Pigs’ brain responses to stroking after long-term positive human interactions: An fMRI exploratory study

**DOI:** 10.1101/2025.08.21.671579

**Authors:** Oceane Schmitt, Yann Serrand, Bibiane Pollak, Giorgio Mattaliano, Nicolas Coquery, Pierre-Antoine Eliat, David Val-Laillet, Jean-Loup Rault

## Abstract

Positive interactions with humans can induce pleasurable experiences in animals, but their underlying neurobiology mechanisms are unknown. We investigated the brain responses (Functional Magnetic Resonance Imaging) to stroking by a human in 20 pigs under general anaesthesia. Ten pigs received positive human contacts over 9 weeks post-weaning (POS) and 10 pigs did not (CTL). Images from CTL pigs showed peaks of activation in the primary somatosensory cortex, caudate nucleus, anterior prefrontal cortex, and dorsal anterior cingulate cortex during stroking. Greater peaks of activation in the anterior prefrontal, ventral anterior cingulate, primary somatosensory and somatosensory association cortices were observed in POS pigs; whereas greater peaks of activation in the amygdala were observed in CTL pigs. Therefore, stroking is perceived and possibly elicited positive emotions in anesthetised pigs, and it may be perceived as more pleasant by experienced pigs than naive pigs, which may rather perceive it as a novel stimulus.

**Highlights:**

- The brain of anaesthetised pigs can perceive tactile stimuli
- Stroking by humans is a pleasurable experience for pigs
- Experience with positive contacts enhances signs of positive perception of stroking

## Introduction

Human-animal interactions form an inherent part of the daily life of domestic animals and can have profound impact on animal welfare^1,2^. A vast amount of scientific evidence demonstrates that negative human-animal interactions (*e*.*g*. loud noises, sudden movements, pushing, hitting and forceful contact) increase the animals’ fear of humans and trigger the stress response in presence of humans in general^3^. More recently, studies have investigated positive human-animal interactions (*i*.*e*. involving soft voice, calm movements, gentle strokes and scratches, and letting the animal choose to approach or not) and found that they bring benefits to animals^2,4^. For instance, pigs stroked 5 min per day during lactation showed better resilience (*i*.*e*. reduced stress response) to handling^5^ and social isolation^6^. Beyond reducing fear of humans and increasing productivity^7^, positive interactions can lead to the establishment of a positive human-animal relationship and trigger indicators of positive emotional states in animals (*e*.*g*. pigs^8^ and rats^9^). Human-animal interactions often involve gentle tactile contacts (*i*.*e*. strokes, scratches) that are generally characterized as positive based on the behavioral responses of the animal, which seeks and maintains the contact and might be in a relaxed or confident state (*e*.*g*. closing eyes, exposing belly). To a certain extent, stroking imitates allogrooming, a positive behaviour occuring between prefered social partners which consists of gently touching, licking, rubbing, nibbling or sniffing another pig ^10,11^, although it has its own pattern and sensory properties (*i*.*e*. repeadly applying the hand on the body of the animal). Domestic pigs naturally seek contact and proximity with humans, and enjoy being stroked^12,13^. However, it is still unclear how these tactile interactions are processed by the animal at the brain level, and which neurocognitive mechanisms involved.

The first brain response to stroking is expected in the somatosensory cortex, which processes tactile stimulations. For instance, tickling rats results in increased firing of cells in the somatosensory cortex^14^. Furthermore, positive tactile interactions are also expected to constitute social-like (*e*.*g*. release of oxytocin during or after interactions^15^) and rewarding situations (*e*.*g*. heifers pursuing humans for strokes^16^), associated with pleasure (*e*.*g*. pigs relaxing while stroked^17^), and would thus possibly mobilize brain regions involved in these processes.

Reward learning consolidation is well known to involve the striatum (basal ganglia) together with associative and integrative brain structures such as the amygdala, hippocampus, prefrontal cortex, and anterior cingulate cortex^18^. For instance, sucrose administration in pigs, which represents a palatable gustatory stimulus, activated the right putamen and right caudate, but also the (anterior and dorsolateral) prefrontal cortex and the anterior cingulate cortex^19^. However, pleasant stimulations are translated into different neuronal patterns of activation depending on their nature and sensory modality^18^. For instance, access to a social reward (*i*.*e*. images of conspecifics) activated the caudate neurons whereas access to a juice reward activated the putamen and ventral striatum neurons in macaques^20^. Pleasant touch resulted in the activation in several areas such as the putamen and caudate^21^, pregenual anterior cingulate cortex^22^, the prefrontal cortex^23^ and the anterior cingulate cortex^24^ in humans. More specifically, pleasantness ratings of touch predicted activation in the anterior cingulate cortex^25^. A specific region of the insular cortex plays a central role in interoception and is involved in the processing of affective touch through the C-tactile (CT) afferent nerves. Indeed, stroke lesions in the insula disrupted the perception of affective touch in humans^26^. Furthermore, people being stroked at the optimal speed for eliciting CT afferents discharge (3 cm/s) showed a greater BOLD response in the posterior insula than people stroked at non-optimal speed (30 cm/s)^27^. Functional connectivity analyses showed a coactivation in the medial prefrontal cortex and the dorsal anterior cingulate cortex with the left insula and amygdala during tactile stimulation (*i*.*e*. arm touch), which demonstrated the existence of a network of brain regions beyond the insula involved in coding CT-targeted affective touch^28^. Given the similarities between the pig and human skins^29^ (both having tactile C-fibers) and brains^30^, it can be expected to observe a reaction to tactile stimulation similar to those observed in humans. However, many studies on the perception of pleasant stroking were performed in conscious humans or animals, and it is unsure how well their results would translate to anaesthetized animals. Functional MRI (fMRI) studies in pigs showed that their brain still respond to olfactory^31–34^ or gustatory^35^ stimulations of different valence, and that the hedonic responses to food sensory stimuli depended on previous individual exposures and associative conditioning ^32,35^.

Being touched is not only about the pleasure experienced, but also about the social (or pseudo-social in the case of human-animal interactions) significance. In humans, brain activity measured using functional near-infrared spectroscopy (fNIRS) revealed that the activity in the prefrontal cortex was higher when petting a dog compared to petting a plush animal, especially after repeated exposures^36^. Neophobic reactions to unknown stimulations, even if they are intended to be positive in nature, are translated in the brain by the activation of brain areas involved in alertness and vigilance. For instance, non-social tactile stimulation in the form of gentle airflow applied on the face elicited a greater response of macaques’ amygdala neurons compared to facial grooming-like finger sweeps by a trusted human^37^. The caudate also seems to be involved in neophobic reactions, as less activation in the caudate was observed after olfactory exposure in naive but not experimented anaesthetised pigs regarding a specific odour^31^.

The present study investigated the functional magnetic resonance imaging (fMRI) BOLD responses of anaesthetised pigs to stroking on the back by a human, and their modulation based on previous experience. Two experimental treatments were compared: Pigs that experienced receiving regular positive human contacts (POS) and pigs that did not (control, CTL). We expected to observe a similar response to stroking in the primary somatosensory cortex in both treatments, as a confirmation that the tactile stimulation was perceived and processed by the brain even under anesthesia. We expected to observe a greater response in the prefrontal cortex, in the dorsal striatum (caudate and putamen), in the nucleus accumbens, and in the anterior cingulate cortex in POS pigs compared to CTL, and a greater activation in the amygdala in CTL pigs compared to POS.

## Results

### Brain response to touch

Touch-elicited brain activation were detected in the right caudate nucleus, the right (two clusters) and left anterior prefrontal cortex, the right primary somatosensory cortex, and right dorsal anterior cingulate cortex as seen in the brain activation map of naive, control pigs (**Figure 1**), which was confirmed by the corrected SVC-based statistics (**Table 1**).

**Table 1.**
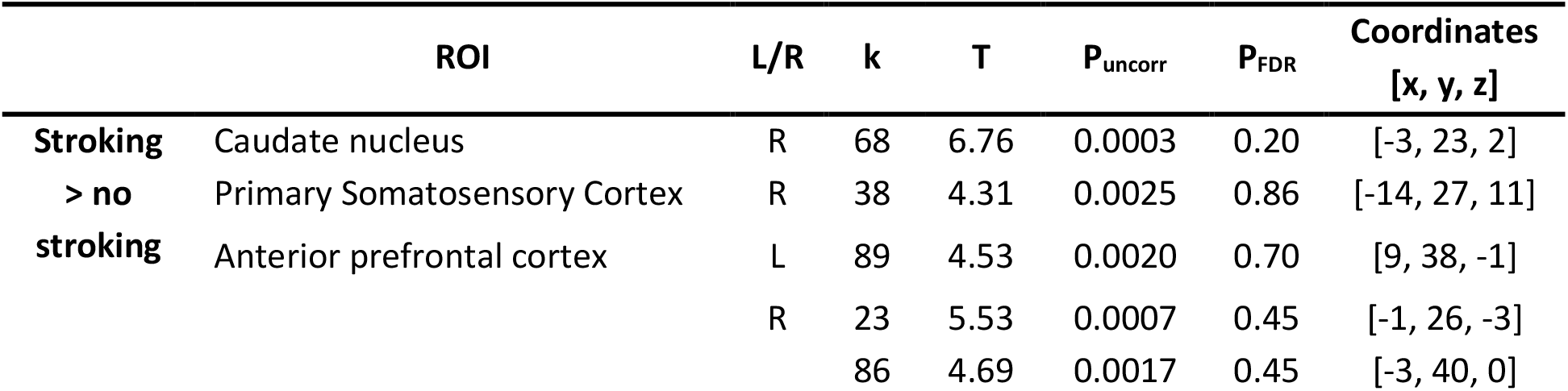
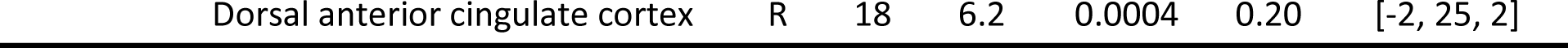
SVC-based statistics showing the brain responses to tactile stimulation (*i*.*e*. comparing stroking bouts to no stroking bouts) of pigs (n=8) that were naive to human stroking (control). Related regions of interest (ROIs) with uncorrected *p*-value (P_uncorr_) that reached the criteria of *p* < 0.05 after Bonferroni correction (ROI number = 18). The test statistic (T) and the corrected *p*-value (False Discovery Rate, P_FDR_) are also presented, along with the laterality of the ROI (L = left, R = right), the size (k) and the coordinates of the cluster. Coordinates were calculated using the pig’s brain atlas^38^.

**Figure 1.**
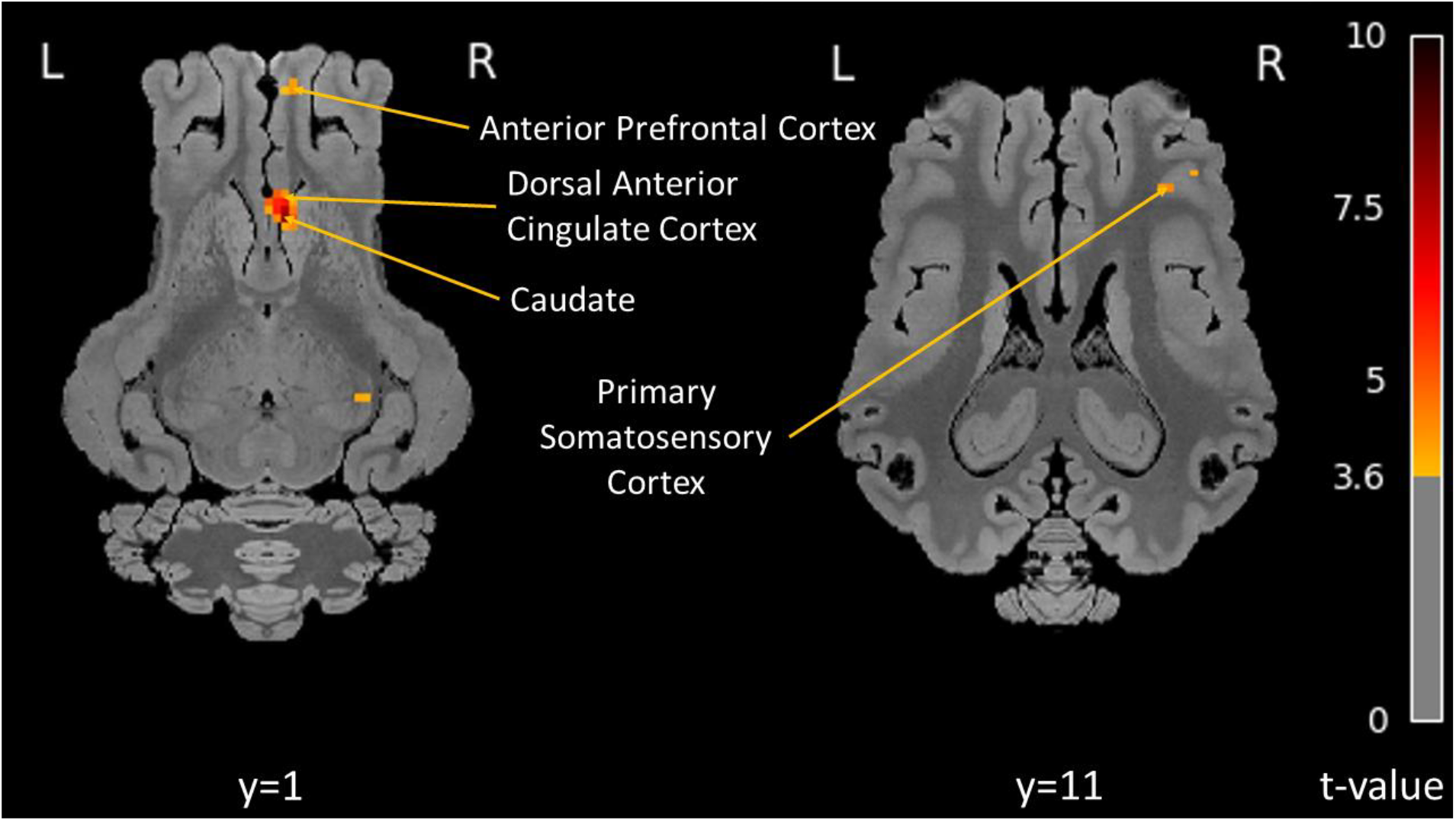
Brain activation maps in response to tactile stimulation (stroking) in control pigs (n=8), naive to human stroking (p_FDR_ < 0.05; number of voxels per cluster k > 18). Horizontal maps of brain BOLD responses are represented at different dorsal-ventral levels (y) related to the posterior commissure (in mm). T-values are represented by the color scale on the right. L = left; R = right.

### Effect of positive human contact on the brain responses to tactile stimulation (stroking)

The brain activation maps (**Figure 2**) and the corrected SVC-based statistic (**Table 2**) showed a difference in brain responses to touch between POS pigs and CTL pigs. Control pigs showed greater activation in the right amygdala than POS pigs, whereas POS pigs showed greater activation in the right anterior prefrontal cortex, the right ventral anterior cingulate cortex, the left somatosensory association cortex and the right primary somatosensory cortex compared to CTL pigs. There was no difference between the two treatment groups in the brain activation of nucleus accumbens, caudate or putamen.

**Table 2.**
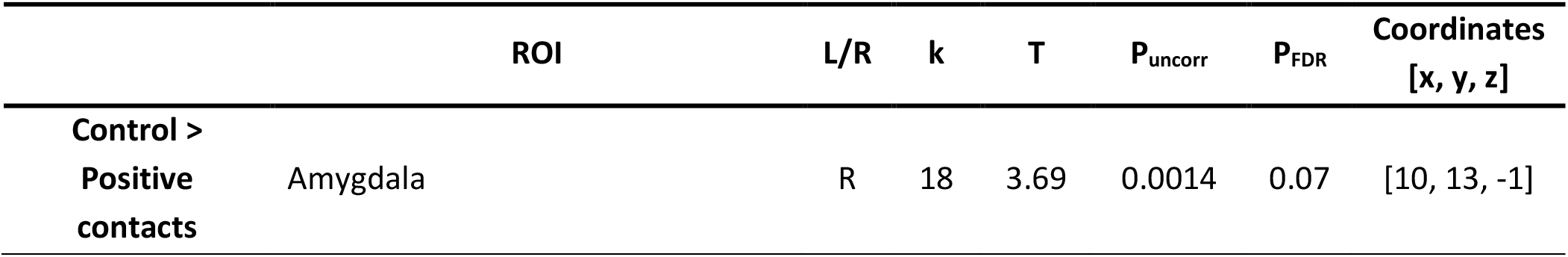
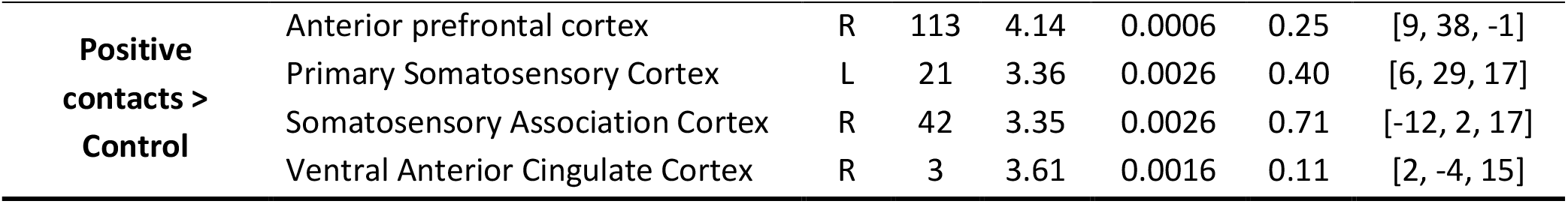
SVC-based statistics comparing the brain response to tactile stimulation (stroking) of the control pigs (n = 8) and the pigs that had received regular positive human contacts (*i*.*e*. strokes; n = 7). Related regions of interest (ROIs) with uncorrected p-value (P_uncorr_) that reached the criteria of p < 0.05 after Bonferroni correction (ROI number = 18). The test statistic (T) and the corrected p-value (False Discovery Rate, P_FDR_) are also presented, along with the laterality of the ROI (L = left, R = right), the size (k) and the coordinates of the cluster. Coordinates were calculated using the pig’s brain atlas ^38^.

**Figure 2.**
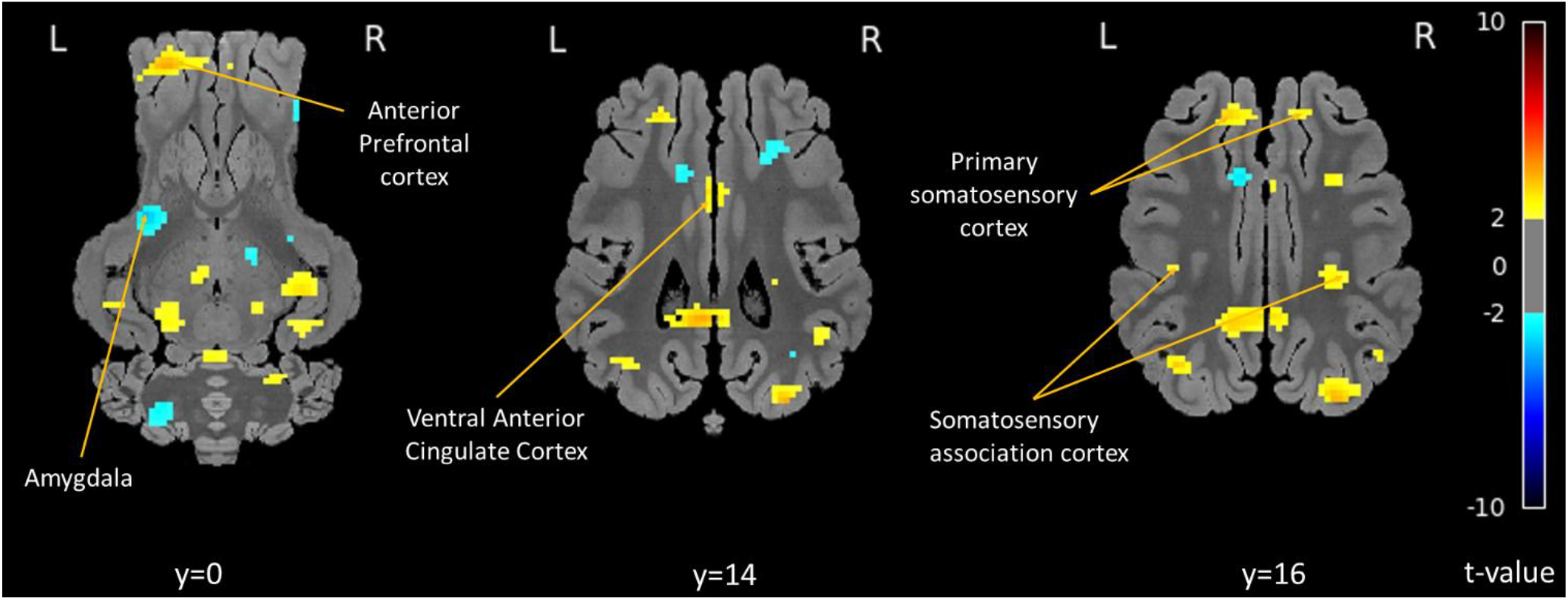
Contrast maps showing the peaks of activation in response to tactile stimulation in the brain of control pigs (in blue) and pigs which received positive human contacts (in yellow) (p_uncorr_ < 0.05; number of voxels per cluster k > 3). Horizontal maps of brain BOLD responses are represented at different dorsal-ventral levels (y) related to the posterior commissure (in mm). T-values are represented by the color scale on the right. L = left; R = right.

## Discussion

Touch by a human in the form of stroking elicited brain responses in pigs, even under general anaesthesia, with activation in the primary somatosensory cortex, but also in the caudate nucleus, anterior prefrontal cortex, and dorsal anterior cingulate cortex. These results suggest that the pigs perceived the tactile stimuli, and that gentle stroking by a human was perceived as pleasant. In addition, pigs experienced with receiving regular positive human interactions showed greater activation in the prefrontal, ventral anterior cingulate, primary somatosensory and somatosensory association cortices, suggesting that stroking was perceived as more pleasant or at least that it mobilised integrative and higher-cognitive brain processes, possibly involved in positive emotions. In contrast, control pigs showed greater activation in the amygdala, possibly due to the novelty of the situation for them.

The results of this study first confirmed that pigs under general anaesthesia perceived tactile stimuli, in the form of a gentle stroke along their spine. Other studies found that pigs under anaesthesia could perceive olfactory^31–34^, gustatory^19,35^ or olfactogustatory (*i*.*e*. flavour)^39,40^ stimulations, as evidenced by differences in brain responses (BOLD with fMRI, blood brain perfusion with SPECT or glucose metabolism with PET) to the stimuli according to their previous experiences. In the present study, we compared the BOLD responses of the pig’s brain with and without stimulation, using CTL pigs to assess the naive response to the tactile stimulation. Brain activation differences were observed in some of our *a priori* brain regions of interest (ROI): The right caudate nucleus, the right and left anterior prefrontal cortex, the right primary somatosensory cortex, and right dorsal anterior cingulate cortex. This is in accordance to the human literature reporting that pleasant touch resulted in the activation of the caudate^21^, anterior cingulate cortex^22,24^ and prefrontal cortex^23^. Furthermore, in macaques, affective touch activated the insula and the anterior cingulate cortex^41^. Similar responses (*i*.*e*. activation in the right caudate, anterior prefrontal cortex, ventral anterior cingulate cortex) were observed in anesthetised pigs that received paired oral and duodenal sucrose infusion compared to pigs that received only saline treatment, which was interpreted as an hedonic response^19^. Therefore, our results suggest that the stimulation was perceived as pleasurable by the pigs, even in those naive to human contacts. However, contrary to our expectations based on the human literature reporting that stimulation of C-tactile afferents leads to an activation in the insula^42^, we found no difference in this area. This could be due to a failure to meet the optimal stroking velocity (3 cm/s in humans and non-human primates) and/or pressure for the stimulation of C-tactile afferents^43^. Indeed, greater activation of the insula was observed in the brain of lightly anaesthetized macaques when they were stroked at 3 cm/s compared to when they were stroked at 15 cm/s^41^. Moreover, while the insula is preferentially activated by touch of low velocity, the primary somatosensory cortex is specifically and preferentially activated by touch of higher velocity (*i*.*e*. discriminative touch) (Morrison, 2016), which is also in accordance to our results. While not matching the affective touch velocity, the tactile stimulation (stroking at approximately 20 cm/s) during the MRI scan was still not as fast and dynamic as the positive contacts provided during the treatment period, which the pigs seemed to enjoy based on their behavioural response (*i*.*e*. maintained contacts, pressed body against the hand, grunted and rounded their backs). Therefore, further studies could also investigate the perception of more rapid strokes and gentle scratching.

The second major finding of our study is the influence of long-term exposure to positive contacts with a human on the pigs’ brain responses to stroking. Pigs experienced to receiving regular human contacts showed activation differences in some of our pre-defined regions of interest compared to control pigs during the tactile stimulation. In agreement with our predictions, the BOLD response of POS pigs was higher in the left ventral anterior cingulate cortex, right anterior pre-frontal cortex, while CTL pigs had greater activation in the left amygdala. However, contrary to our predictions, there were no activation differences between groups in the putamen, caudate and insular cortex. Unexpectedly, there was a higher BOLD response in the primary somatosensory cortex and somatosensory association cortex in the POS pigs.

In humans, the perception of the pleasantness of a tactile stimulation seems to be influenced by the level of exposure to touch. Indeed, humans with low touch exposure (weekly or monthly) rated the pleasantness of being touched at the optimal velocity (3 cm/s) significantly lower than humans with a higher touch exposure (daily or weekly)^45^. For the other (suboptimal) velocities, both groups rated them as less pleasant and did not differ in their ratings^45^. This supports our finding that control pigs, although perceiving touch as pleasant, perceived it as less pleasant as pigs who received this contact three times weekly over nine weeks. However, the question remains whether pigs that received more allogrooming from their counterparts would react differently to stroking by a human. Allogrooming would result in a different sensory experience since it involves gentle nibbling and licking, thus leading to saliva deposits and calorific exchanges due to the variations of humidity on the skin, which is not the case with a human hand stroke. Further studies are warranted to elucidate the effect of intra-species touch on the perception of inter-species touch, and the extend of the inter-individuality variability in the perception of (pleasant) inter- and intra-species touch.

It has been proposed that the anterior cingulate cortex is the centre for action-output learning, implying its involvement in emotional process and memory of the link between rewards and actions^46^, but also in more complex processing such as personal meaning in humans^47^ (i.e. it is activated by memories of intrinsically motivated events^48^). The prefrontal cortex also plays a crucial role in processing, integrating and regulating experiences, including social interactions^49^. Specifically, the anterior prefrontal cortex has been hypothesised to play a role in recollection of contextual “internally generated” information (*e*.*g*. emotions) related to a past event (*e*.*g*. in our study, the positive contacts received in the home pen)^50^. In the context of the present study, those findings in humans suggest that the tactile stimulation triggered neural mechanisms linked to information recollection and emotional regulation in the POS pigs, possibly showing that they associated the stroking stimulation with their past experience with the human in the home pen and that it may carry a personal meaning for those pigs. However, since we could not find significant differences of activation in the caudate and putamen, which are also involved in reward processing^18^, our results suggest that stroking along the spine was marginally more pleasant in POS pigs, compared to CTL pigs.

The differences in the activation in the amygdala were the most significant of all regions considered in the analysis. Our hypothesis was that we would observe a greater activation in the CTL pigs, for which the tactile stimulation from a human was an unknown stimulus, as a marker of surprise. Indeed, while both hippocampus and amygdala respond to novelty, only the amygdala was observed to have a greater BOLD response to unusual (*vs*. common) novel visual stimuli in humans^51^. Several studies found higher activation in relation to pleasant touch (*e*.*g*.^52^). However, in the context of our results, it would seem more intuitive to interpret the greater activation in the amygdala in the CTL pigs as a marker of surprise or novelty, given that stroking was a novel type of experience for them, compared to the POS pigs experienced to interacting with the human in various ways including being stroked on the back.

Finally, the greater activation in the primary somatosensory cortex and in the somatosensory association cortex in the POS pigs were unexpected, as we assumed that its activation was purely due to the processing of the tactile stimulation. For instance, the neurons in the somatosensory cortex have a similar response to the social and non-social tactile stimuli in macaques^37^. However, it is also possible that the experience with positive tactile contacts participated in the development of a greater neuroplasticity in the POS pigs, resulting in more neural activation of the somatosensory cortex during the tactile stimulation^53^. For instance, humans became more sensitive to androstenone (*i*.*e*. lower detection threshold) through repetitive exposure, which was accompanied by an increased neural response (*i*.*e*. increased olfactory evoked potential measured by an electroencephalogram^54^). There is also some disagreement between studies on whether the somatosensory cortex and/or the anterior cingulate cortex encode the affective value of touch, as some studies found that pleasantness ratings of touch predicted activation in the anterior cingulate cortex and not in the somatosensory cortex (*e*.*g*. ^25^), but others found that the somatosensory cortex only was involved (*e*.*g*. ^55^). Therefore, our study might bring further evidence that the somatosensory cortex may indeed be involved in the affective processing of touch.

Collectively, our results suggest that stroking is genuinely perceived as pleasant by pigs, and that experience might make the perception more salient. This is also supported by behavioural data (not shown). Indeed, on the last day of the experiment, after the MRI scans, the voluntary approach to the human was measured for all pigs and they were offered a 10-min positive contacts session in their home pen (*i*.*e*. similar to POS treatment). Despite taking longer to approach the human^56^, CTL pigs spent 36% of their time in contact with the human (POS: 55%, data not shown). However, future studies should use additional measures of emotional response (*e*.*g*. heart rate variability) to further assert our findings. The most obvious limitation of this exploratory study is the low number of pigs involved, which implied a high inter-individual variability. Furthermore, our results have to be taken with caution, especially because our study design implies inferring mental states and processes from the activation of specific brain areas, i.e. reverse inference, which can be misleading because the activated brain areas could be involved in other mental processes than the ones considered^57^. A final word of caution should be added regarding the effects of the anaesthetic products on the BOLD responses. We avoided the use of inhalant anaesthetics (e.g. isoflurane, sevoflurane) because these agents are known to affect neurovascular coupling and cerebral blood flow,leading to reduced neural activity and impairing the quality and interpretation of fMRI^58,59^. Instead, medetomidine was administered to maintain deep sedation. Medetomidine is commonly used in fMRI studies due to its relatively stable effects on cerebral haemodynamics and neural activity, making it a suitable choice for preserving BOLD signal integrity during imaging^60^. Although the effects of azaperone on BOLD response remain unstudied, ketamine has been reported not to significantly alter BOLD signals related to cognitive states^61^.

However, our results are quite encouraging for using larger sample sizes in future studies and represent a princeps demonstration of the neurocognitive effects of positive tactile interactions with humans in pigs.

## Conclusions

Stroking by a human was perceived and processed by the brain of the pigs, even under general anaesthesia. Naive pigs (CTL) seemed to process the stroking stimulation as an unknown or surprising stimulus. Previous experience of regular positive human contacts enhanced the POS pigs’ response to stroking, with activation in brain regions involved in the processing of pleasurable and social stimuli. Therefore, although all pigs seemed to process stroking as a potentially pleasant experience, this perception seemed to be stronger in the experienced pigs. Those results not only confirmed the translation of previous findings between species, but they also provide novel knowledge regarding the perception of human gentle contacts by animals. Finally, this study contributed to further validating neuroimaging as a tool for the assessment of pigs’ positive emotions and welfare, especially in the context of human-animal interactions and physical contacts.

## Materials and methods

The material and methods of the study, as well as the rationale behind the selection of the ROIs, have been submitted as a pre-registration on OSF^62^ and can be found here: https://doi.org/10.17605/OSF.IO/HKR67

### Animals and housing

This study involved 24 female pigs (*Sus Scrofa domesticus;* Swiss Large White × Pietrain breed) used over two batches (6 pigs per group for each batch). Only female pigs were used in this study as a mean to reduce variability and sample size, and because the overall project (that this study is part of) focused on female pigs. Pairs of female siblings were recruited at weaning, based on being healthy and of average weight. One sibling was allocated to the “positive human contacts” (POS) group and the other sibling was allocated to the “control” group (CTL). Pigs were housed in trios, making four trios (two per treatment) per batch. Twenty of these pigs (10 per treatment, all sibling pairs) were randomly selected and scanned in a 1.5T Magnetic Resonance scanner (Siemens Healthineers, Magnetom Espree 1.5 T; Siemens Medical Solutions) once at weaning (before the treatment started, at 5 weeks of age) and the same individuals at the end of the treatment (16 weeks of age).

### Treatments

After the first MRI (Magnetic Resonance Imaging) session, treatments were applied to create a different experience regarding human-animal interactions. In the “positive human contacts” group (POS), pigs got opportunities to interact voluntarily with a human and received positive contacts (such as petting, stroking, scratching) for 10 min, twice a day (10:00-11:00 and 14:00-15:00), three days a week (Monday, Wednesday and Friday), over 9 consecutive weeks. The positive contacts could be applied on any part of the pig’s body, in an individual manner depending on what the pig seemed to enjoy, but mainly occurred on the neck (scratching behind the ears, stroking), back (stroking and scratching) and belly (rubbing). In the “control” group (CTL), pigs were only exposed to the presence of the same human, who was standing silently outside and in front of their home pen and avoiding eye contact, following the same schedule as for POS pigs.

### Anaesthesia protocol

Each pig underwent MRI scanning under general anaesthesia, performed by a certified veterinary anaesthetist (GM), using a refined protocol designed to optimize BOLD image acquisition ^63^. The same anaesthetic protocol was used in all pigs and doses were adjusted according to age group and demeanour in order to minimize stress in the perianaesthetic period. Pigs were fasted for 4-8 hours with access to water ad libitum until premedication. Each pig was sedated intramuscularly with ketamine (Narketan^®^, 7-14 mg/kg Body Weight [BW]) and azaperone (Stresnil^®^, 1-2 mg/kg BW) into the brachiocephalic muscle. Following injection, pigs were left undisturbed in a familiar, dark room (their home enclosure) for a variable period (5-20 minutes) until moderate sedation was achieved. Distance monitoring was used to avoid additional stress. If sedation was deemed insufficient, a second intramuscular injection of one-third to one-fourth of the initial doses was administered before transport to the preparation room. Pigs were kept in sternal recumbency on a padded table. A catheter (20G or 22G) was placed in the external auricular vein and secured. After preoxygenation for at least 5 minutes with 100% oxygen via mask, general anaesthesia was induced with ketamine (Narketan^®^, 20-30 mg/kg BW), azaperone (5 mg/kg BW) and atropine (0.04 mg/kg BW) intravenously. Pigs were orotracheally intubated with a cuffed endotracheal tube of appropriate size and the tube was secured. A second catheter (20G or 22G) was placed in the central auricular artery for invasive blood pressure monitoring. Eye lubricant was applied to prevent corneal drying. Pigs were then transported to the MRI suite, where the endotracheal tube was connected to a circle breathing system in 100% oxygen and to the anaesthetic machine. Mechanical ventilation was initiated, with tidal volume and respiratory rate adjusted to maintain end-tidal CO_2_ between 40 and 45 mmHg. A continuous intravenous infusion of medetomidine (Domitor^®^, 0.018 mg/kg/h) was initiated. An isotonic crystalloid solution (Sterofundin ISO) was administered at a variable rate (5-10 ml/kg/h) throughout the procedure. Standard perioperative monitoring and care was applied and recorded: anaesthetic depth, heart rate, electrocardiography, respiratory rate, pulse oximetry, capnography, spirometry, invasive blood pressure and rectal temperature were monitored using a multi-parameter monitor (HP CMS-2000 Anaesthesia monitor) throughout the procedure. Mean arterial blood pressure, arterial haemoglobin oxygen saturation and body temperature were maintained above 60 mmHg, 95% and 36°C, respectively. At the end of the procedure, pigs were allowed to recover under close supervision. Atipamezole (Antisedan^®^, 0.05-0.1 mg/kg) was administered intramuscularly to antagonise the effects of medetomidine, facilitating recovery. Following tracheal extubation, oxygen was administered via facemask to support recovery. Intravenous crystalloid infusion was continued until the pigs were able to stand unaided. Once stable, each pig was returned to its enclosure and kept separated from the rest of the group until fully awake. The entire procedure, from sedation to end of anaesthesia, lasted up to three hours. Recovery times varied among individuals, and was up to two hours.

### MRI scans procedure

For the need of the scanning procedure, the pig’s head was placed inside a human coil (knee coil at 5 weeks of age, head coil at 15 weeks of age). The scanning session lasted approximately 60 min and the MRI sequence included structural and functional imaging (**Figure 3**). The images were acquired using the software Syngo.via (Siemens Healthcare GmbH, Germany). The details of the acquisition characteristics of each MRI sequence can be found in **Table 3**. The Blood-Oxygen-Level-Dependent (BOLD) functional imaging included a “resting state” (rsMRI), which aimed to assess the general neuronal activity and connectivity of the pigs’ brain, followed by a “stimulation” (BOLD fMRI), which aimed to assess pigs’ neuronal activity in response to stroking. The human involved in the weekly interactions (“familiar human”) performed the tactile stimulation during the fMRI, given that pigs may perceive and discriminate humans based on smell (and maintain a sense of smell even under anaesthesia, *e*.*g*. ^31^). During fMRI, the familiar human entered the MRI room and alternated between non-stimulation (*i*.*e*. just standing next to the scanned pig in silence) for one minute, and stroking the scanned pig on her back (soft stroke made by applying gently the full palm of the right hand, from shoulders to rumps along the spine) for one minute (1 stroke every 3 seconds, timing based on the pulses of the MR scanner) for a total of 20 repetitions. Strokes were delivered from the cranial to the caudal region of the back, hereby following the direction of hair growth. To avoid confounding effects, the interacting human was not involved in the handling of the pigs for the scanning procedure and remained in a separate room until the time of stimulation.

**Table 3.** Details of the acquisition characteristics of each MRI sequence.

|  | TR<br>Repetition<br>time (ms) | TE<br>Echo time (ms) | FOV<br>Field of view<br>(mm × mm) | Matrix size | Number of<br>slices |
| --- | --- | --- | --- | --- | --- |
| T2 | 5370 | 109 | 240x240 | 256x256 | 46 |
| T1_sag | 1170 | 5.58 | 230x230 | 256x208 | 88 |
| T1_tir | 7000 | 63 | 180x180 | 256x256 | 52 |
| FLAIR | 8500 | 86 | 180x180 | 512x448 | 25 |
| DTI | 3200 | 105 | 204x204 | 104x128 | 20 |
| Field Map | 460 | 4.78/9.54 | 192x192 | 64x64 | 36 |
| BOLD (T2*) | 3000 | 39 | 192x192 | 74x74 | 22 |

**Figure 3.**
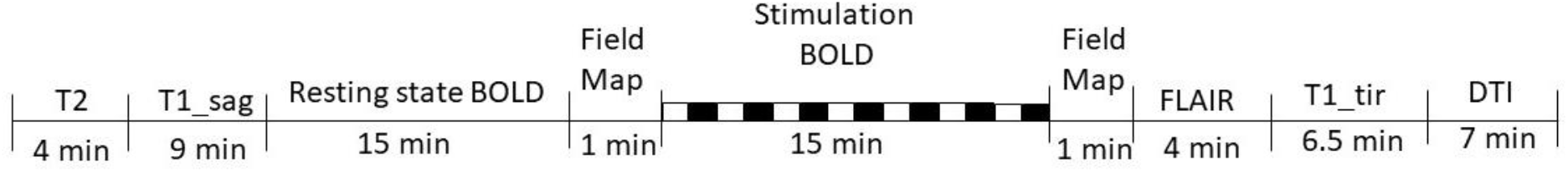
Schematic representation of the MRI session timeline. Within the “Stimulation BOLD” section, the black rectangles represent the 1-min periods of stroking stimulation (*i*.*e*. human stroking the pig’s back), and the white rectangles represent the 1-min periods without stimulation (*i*.*e*. human standing next to the pig in silence). T2: Transverse relaxation time, allows characterization of tissues by capturing differences in decay latency of spins’ aligned precession in the transverse plane. T1_sag: Longitudinal relaxation time performed in transversal view, allows characterization of tissues by capturing differences in the proton spins’ latency to relaxation from the sagittal plane toward the main longitudinal magnetic vector. T1_tir: Longitudinal relaxation time with turbo inversion recovery, similar to FLAIR. FLAIR: Fluid Attenuated Inversion Recovery, removes the signal from the cerebrospinal fluid and therefore allows to detect subtle changes at the periphery of the hemispheres and in the periventricular region close to CSF. DTI: Diffusion Tensor Imaging, allows mapping white matter’s nerve tracts. BOLD: Blood-Oxygen-Level-Dependent imaging, allows quantifying the variations of oxygen transported by haemoglobin in relation to neuronal activity.

### Processing of images

For the present study, only the data (*i*.*e*. volumes representing BOLD signal in voxels) from the images acquired during the stimulation period (fMRI) were analysed, using SPM12. A segmented 3D atlas of the pig brain^38^ allowed to identify the regions of interest (ROIs), and to define the masks used for small-volume correction.

The selected regions of interest (ROIs) where those for which evidence of difference of activation due to social contact and/or pleasurable experience was found in the scientific literature, in pigs or other species (as cited in the introduction):

- Anterior prefrontal cortex
- Anterior cingulate cortex
- Insular cortex
- Dorsal striatum (putamen and caudate)
- Ventral striatum (nucleus accumbens)
- Amygdala
- Somatosensory cortex
- Somatosensory association cortex

Segmentation was performed using SPM12 Segmentation algorithm. Pre-processing of BOLD sequences consisted in slice-timing, realignment, coregistration, normalization and smoothing. All those steps were performed using SPM12 corresponding algorithms and Nipypye for pipelining.

### Statistical analysis

The statistical unit was the individual pig. We used a whole-brain approach, but also applied small-volume correction (SVC) for our *a priori* regions of interest (ROI). Due to limitations related to the size of the pig’s brain and the effect of anaesthesia on brain activity, we used a non-standard statistical analysis with regards to human statistical standards usually considering statistical significance at a cluster level with p-value < 0.05^32^. For the whole brain approach we used a threshold set at p < 0.05 (with a False Discovery Rate (FDR) correction for the effect of touch analysis or without correction for the effect of positive contacts vs. control analysis) to produce the brain activation maps, and for the SVC we used a p-value corrected for multiple ROIs comparisons with a Bonferroni correction and at a threshold of 0.05 (peak level). Nine brain regions were analysed, corresponding to 18 ROIs bilaterally, thus the correction was applied for n = 18 ROIs. Corrected and uncorrected p-values are presented in the final results, since our study is exploratory and performed on a low number of animals^32,35^.

To investigate the differences in the brain responses to tactile stimulation (comparing images of naive pigs with and without stimulation), we ran a first-level analysis on each pig using a mass-univariate approach based on General Linear Model (GLM). Betas estimators of the model were then used to compute the different contrasts. Those contrasts fed the second-level analysis, also referred as “Random-Effects” or “Mixed-Effects” analysis as it considers both within- and between-subject variance. For this hypothesis, a one-sample t-test was used to compare brain responses to tactile stimulation and non-stimulation bouts in CTL pigs (n=8) at the second time point (“after” treatment application). The random effects of pig, pen and replicate were accounted for in the models.

To investigate the difference due to exposure to positive human contacts, we ran a two-sample t-test-factor ANOVA, where we compared pigs from the two experimental treatments (CTL n = 8; POS n = 7) at the second session (“after” treatment application). The random effects of pig, pen and replicate were accounted for in the models. We looked at activation differences using both a whole-brain and a SVC approach using the same procedure as above.

Some data were excluded from the analysis, due to the following reasons:

- Whole sequence, problems due to acquisition:
  - misplacing of the pig (n = 2, pigs 8485 and 9391)
  - intensity problem (n = 1; pig 8599)
  - artefact in the sequence in the prefrontal region (n = 2; pigs 8514 and 9349)
- Volumes in sequence (n = 122 volumes excluded from 30 sequences; range: 0-19 volumes excluded per sequence):
  - volumes with too much displacement (rotation and/or translation) compared to the preceding ones
  - volumes with too much change in intensity compared to the preceding ones

## Data Availability Statement

All data generated or analysed during this study are included in this article. The dataset containing the results of the statistical analyses can be found on Open Science Framework repository: https://osf.io/b7jad/?view_only=e7d2eaf3a64646d08655590b18421640. Further inquiries can be directed to the corresponding authors.

## Acknowledgements

The author disclose support for the research and Open Access publication of this work from the Austrian Science Fund (Fonds zur Förderung der Wissenschaftlichen Forschung, FWF; grant number P33669-B).

We would like to thank the staff at the Medau farm and at the Swine Clinic of the University of Veterinary Medicine, Vienna for taking care of the pigs and providing help in the conduction of this experiment, and Jason Yee for developing the MRI sequence.

## Authors Contribution

Conceptualisation: OS, JLR; Data curation: YS; Formal analysis: YS, NC; Funding acquisition: JLR; Investigation: OS, BP, GM; Methodology: OS, JLR; Project administration: OS; Resources: JLR, BP, GM, DVL; Visualisation: YS; Writing – original draft preparation: OS; Writing - Review and editing manuscript: OS, JLR, NC, BP, GM, DVL, PAE.

## Ethics declarations

This experiment was conducted in accordance with the national legislation and the European Directive 2010/63/EU. It was approved by the BMBWF (Bundesministerium für Bildung, Wissenschaft und Forschung) under the approval number 2022-0.118.102.

